# Androgen depletion increases sensitivity to effort-related costs and alters mesoaccumbal circuit function in male mice

**DOI:** 10.64898/2026.05.29.728788

**Authors:** Sara R. Westbrook, Qing Wang, Allison L. Jensen, Courtney M. Klappenbach, Katy Touretsky, Kristen M. Delevich

**Affiliations:** Department of Integrative Physiology and Neuroscience, Washington State University, Pullman, WA, USA; Center for Reproductive Biology, Washington State University, Pullman, WA, USA

## Abstract

**Background:** Androgen deficiency in males is associated with reduced motivation, fatigue, and decreased goal-directed behavior, yet the neural mechanisms underlying these changes remain poorly understood. Dopamine signaling within the nucleus accumbens (NAc) plays a central role in regulating effort-based decision making. Here, we tested the hypothesis that loss of testicular hormones alters mesoaccumbal dopamine function to increase sensitivity to effort-related costs.

**Methods:** Male mice underwent orchiectomy (ORX) either before puberty onset or in adulthood. Effort-based decision making was assessed using a progressive ratio 1 closed economy (PR1-CE) task. Dopamine-related function was assessed using systemic haloperidol administration and high-performance liquid chromatography to measure dopamine and metabolites, while whole-cell recordings were used to assess intrinsic excitability of NAc spiny projection neurons (SPNs).

**Results:** ORX increased sensitivity to effort costs, reflected by a shift toward energy-efficient responding while maintaining overall food intake. These behavioral changes were accompanied by reduced responsiveness to haloperidol. Postpubertal ORX increased dopamine content and reduced metabolite-to-dopamine ratios in the NAc, consistent with reduced dopamine turnover, whereas prepubertal ORX did not affect dopamine measures. Prepubertal ORX selectively reduced excitability of NAc core D1R^+^ SPNs, while postpubertal ORX increased excitability across both D1R^+^ and D1R^−^ populations.

**Conclusions:** Androgen depletion increases effort cost sensitivity and is associated with alterations in mesoaccumbal circuit function. Although behavioral effects were similar following pre- or postpubertal ORX, distinct neurochemical and cellular adaptations were observed, suggesting developmental timing influences neural adaptations to androgen depletion. These findings provide insight into neural mechanisms linking androgen deficiency to motivational deficits.

## Introduction

Androgen deficiency in adult males is frequently accompanied by diminished motivational drive/apathy, fatigue, reduced physical activity, and depressive symptoms (Bhasin et al., 2010; Boeri et al., 2025; Shores et al., 2004). Although both age-related and pathological hypogonadism (including iatrogenic forms such as androgen deprivation therapy) produce well-characterized peripheral and metabolic changes (Braga-Basaria et al., 2006; Miller et al., 2024; Muller et al., 2005), the neural mechanisms contributing to diminished motivation remain poorly understood. Although surgical removal of the testes (orchiectomy; ORX) reduces spontaneous locomotion in male rodents (Daan et al., 1975; Ibebunjo et al., 2011; Jardí et al., 2018; Karatsoreos et al., 2007; Kim et al., 2021; Zhang et al., 2011), these findings do not distinguish between a global behavioral suppression and increased sensitivity to energetic costs. Consistent with the latter interpretation, we previously found that ORX reduced food intake only when an operant response (i.e., effort) was required to obtain it (Klappenbach et al., 2023), suggesting that androgen loss alters neural systems involved in effort-based decision making. Supraphysiological doses of testosterone are known to reduce effort discounting in operant tasks (Donovan and Wood, 2022; Wallin et al., 2015), further supporting a role for androgens in regulating cost sensitivity.

The nucleus accumbens (NAc) is a key interface between motivation and action, integrating glutamatergic inputs from frontal cortical regions with dopaminergic inputs from ventral tegmental area (VTA) (Floresco, 2015; Sesack and Grace, 2010). Dopamine (DA) signaling within the NAc regulates behavioral activation and energy expenditure, influencing both operant responding and locomotor output (Cagniard et al., 2006; Di Chiara and Imperato, 1988; Roitman et al., 2004; Salamone et al., 2016, 2007). Critically, NAc DA appears to be particularly important for obtaining food under high-effort but not low-effort conditions (Aberman and Salamone, 1999; Koch et al., 2000; Nicola, 2010; Salamone and Correa, 2002). More recent studies demonstrate that mesoaccumbal DA dynamically tracks the expected value of future reward through state-like signals linked to motivational engagement, governing whether work is initiated and sustained (Berke, 2018; de Jong et al., 2024; Hamid et al., 2016; Mohebi et al., 2019). Activity of NAc core dopamine type 1 receptor (D1R)- and dopamine type 2 receptor (D2R)-expressing spiny projection neurons (SPNs) regulate physical activity and effort allocation for food (Matikainen-Ankney et al., 2023; Walle et al., 2024). Importantly, there are multiple lines of evidence that androgens can act on target cells within the mesoaccumbal circuit by binding to androgen receptor (AR) or estrogen receptor subtypes (ER) (following aromatization of testosterone to estradiol) expressed within VTA and NAc (Almey et al., 2022; Dart et al., 2024; Krentzel et al., 2021; Kritzer, 1997; Low et al., 2017; Simerly et al., 1990; Tobiansky et al., 2018a). Together, these findings suggest that androgen signaling within mesoaccumbal circuits may influence effort-based decision making by modulating DA-dependent processes that govern behavioral energy allocation.

In the current study, we investigated whether the behavioral phenotype caused by ORX can be interpreted through the lens of the “thrift hypothesis” of DA, which posits that shifts in DA function regulate behavioral energy allocation, influencing both voluntary activity and willingness to expend effort for food reward (Beeler et al., 2012a; Beeler and Mourra, 2018). We further compared the behavioral consequences of ORX performed before puberty onset with those of ORX in adulthood to determine whether prepubertal androgen loss produces enduring organizational effects on motivational circuits that differ from the activational effects of androgen depletion in adulthood.

## Methods and Materials

### Animals and Surgery

The current study used 95 male C57BL/6 mice (Exp. 1: 48 mice; Exp. 2: 47 mice) and 31 transgenic B6.Cg-Tg(Drd1a-tdTomato)6Calak/J ((Ade et al., 2011); JAX stock #016204) male mice (Exp. 3) generated in-house from breeders originally obtained from Charles River and Jackson Laboratory, respectively. Mice were weaned on postnatal day (P) 21, group-housed, and maintained on a reverse 12-hour light/dark cycle (lights off at 0700). Mice received prepubertal (P25) or postpubertal (P90) orchiectomies (ORX) or sham surgeries as previously described (Klappenbach et al., 2023). For Exp. 1, mice were individually housed at the start of home cage behavioral monitoring (i.e., 35 days after surgery) with *ad libitum* access to water. For Exp. 2 and 3, mice were group-housed throughout the experiment and had *ad libitum* access to chow (Purina 5001) and water. Procedures were approved by the Institutional Animal Care and Use Committee at Washington State University and followed AAALAC and NIH guidelines.

### Exp. 1: Behavior

The experimental timeline is provided in Figure 1A. 35 days after surgery, mice were trained in their home cage to nose poke for 20 mg grain pellets (Test Diet 5TUM) using Feeding Experimentation Devices (FED3; Matikainen-Ankney et al., 2021). Behavioral testing began at approximately P70 or P135 for the prepubertal and postpubertal surgery groups, respectively, and included assessment of pellet intake under no cost (free feeding) and escalating cost conditions (progressive ratio 1 closed economy, PR1-CE), home-cage locomotor activity using Pallidus activity sensors (Usiyevich et al., 2025), and sucrose preference tests (SPT). Behavioral outcomes were measured at baseline and in response to vehicle (0.3% tartaric acid) or haloperidol (0.25 or 0.5 mg/kg i.p.; Sigma-Aldrich H1512) challenge. At the end of behavioral testing, mice were euthanized with CO_2_, and seminal vesicles were dissected and weighed. More details are provided in the supplemental methods.

**Figure 1.**
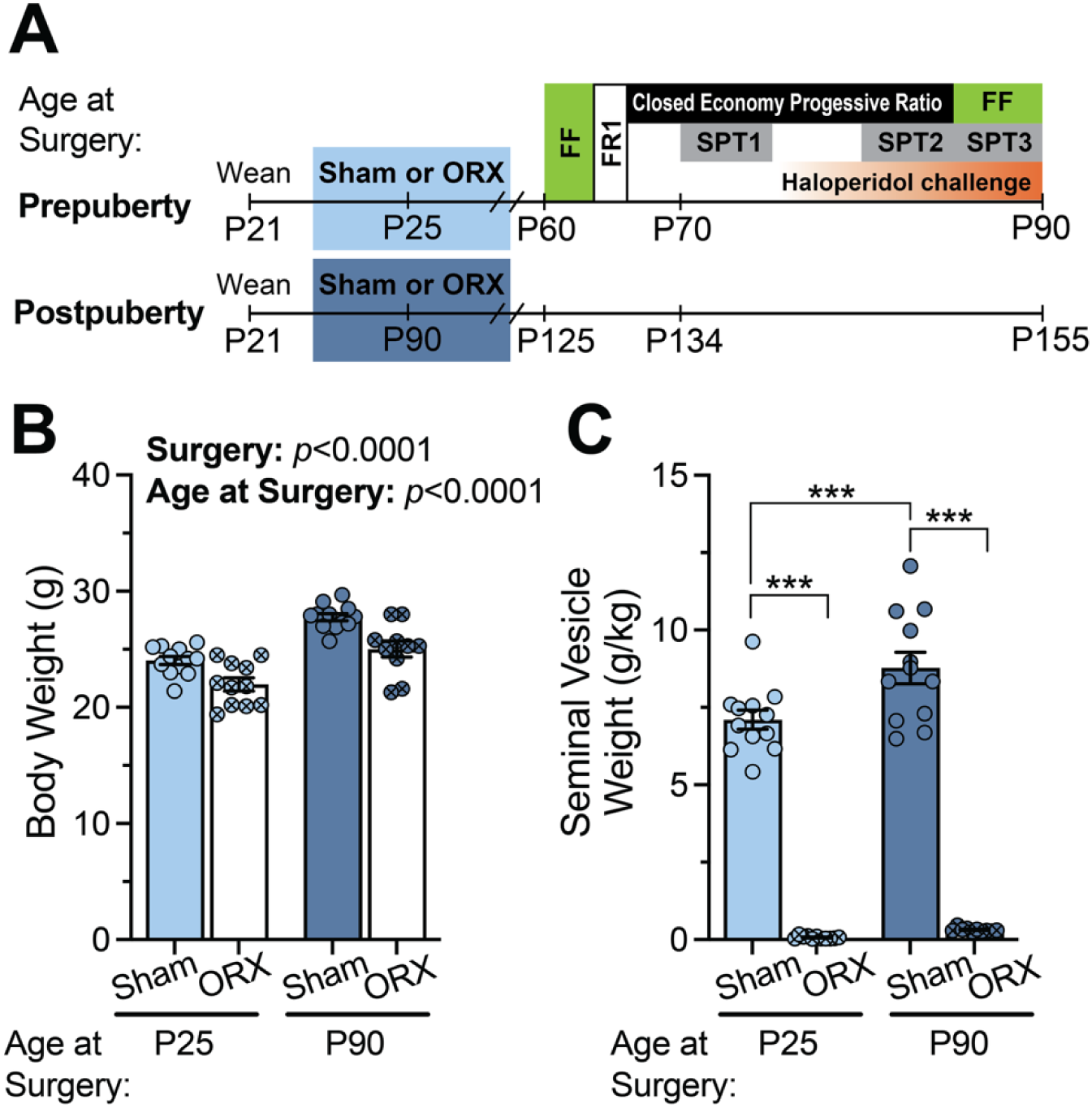
Experimental timeline and ORX validation. (**A**) Experimental timeline for behavioral testing of adult male mice after prepubertal or postpubertal ORX or sham surgery. FF: free feeding; FR1: fixed ratio 1; SPT: sucrose preference testing. (**B**) Terminal body weights (g) at the time of tissue collection. Sham and older mice weighed more than ORX and younger mice (surgery: p<0.0001; age at surgery: p<0.0001, respectively). (**C**) Seminal vesicle weights (g/kg body weight). ORX reduced seminal vesicle weights in both age groups (surgery x age at surgery: p<0.05), indicating successful reduction of circulating testosterone. ***p<0.001, simple main effects

### Exp. 2: Dopamine Content

As shown in Figure 4A, C57BL/6 mice that underwent prepubertal (P25) surgery were euthanized on P70 and those that underwent postpubertal (P90) surgery were euthanized on P135. Briefly, mice were rapidly decapitated, and brains were placed in a matrix and sectioned into 1 mm coronal slices. Sections were maintained on dry ice during tissue collection, and 1 mm^3^ punches were collected from the nucleus accumbens and dorsomedial striatum. Punches from both hemispheres were pooled within each mouse, snap frozen in liquid nitrogen, and stored at −80°C until shipment to Vanderbilt University’s Neurochemistry Core Laboratory for quantification of dopamine and its metabolites using high-performance liquid chromatography with electrochemical detection. More information is provided in the supplemental methods.

### Exp. 3: Electrophysiology

Transgenic Drd1tdTomato mice that received prepubertal (P25) surgery were sacrificed on P60-75 and those with postpubertal (P90) surgery were sacrificed on P125-140, and brains extracted for slice electrophysiology recordings as previously described (Delevich et al., 2022, 2020b). Whole-cell current clamp recordings were conducted in the presence of synaptic blockers (NBQX, DL-AP5, and GABAzine; 10 μM each) to assess intrinsic excitability of Drd1tdTomato+ and Drd1tdTomato- cells in the nucleus accumbens core. See the supplemental methods for more information.

### Data Analysis

Time series data from FED3 devices were processed in R. GraphPad Prism software (version 10, San Diego, CA, USA) was used for graphing, and data are presented as mean ± SEM. Statistical analyses were performed with SAS OnDemand for Academics: Studio statistical software. See the supplemental methods for details.

## Results

To investigate the role of testicular hormones at peripuberty versus adulthood on reward pursuit, male C57BL/6 mice underwent ORX or sham surgery at either prepubertal (P25) or postpubertal (P90) time points. Behavioral testing began 35 days post-surgery (Figure 1A). Sham mice weighed more than ORX mice (main effect of Surgery: F_1,41_=24.35, p<0.0001), with an additional main effect of age at surgery (F_1,41_=48.50, p<0.0001), but no interaction, indicating similar weight reductions following pre- and postpubertal ORX (Figure 1B). ORX significantly reduced seminal vesicle weight in both age groups (Surgery × Age interaction: F_1,41_=4.98, p=0.0312), confirming effective androgen depletion (Figure 1C).

### Progressive Ratio Closed Economy Task

To assess effort-based reward pursuit under baseline or drug challenge conditions, mice were tested in a PR1-CE task in which all food was earned via nose pokes with linearly increasing effort requirements (Figure 2A,B). During baseline, sham mice exhibited greater operant responding than ORX mice, making more active pokes (main effect of Surgery: F_1,40_=20.53, p<0.0001; Figure 2C) and paying higher cost/pellet (i.e., pokes/pellet) (F_1,40_=26.52, p<0.0001; Figure 2D). ORX mice instead adopted a lower-cost strategy, characterized by more frequent ratio resets (F_1,40_=30.14, p<0.0001; Figure 2E). Despite these differences in response strategy, pellets earned did not differ between groups (p>0.05; Figure 2F). Together, these findings suggest increased sensitivity to effort costs rather than reduced appetite. In parallel, sham mice displayed higher home-cage activity levels than ORX mice (F_1,40_=15.75, p=0.0003; Figure 2G). No effects of age at surgery or interactions were observed during baseline. When we examined body weight across 17 days of PR1-CE testing, we observed a main effect of Day (F_5.13,213.9_=7.06, *p*<0.0001), driven by a decrease in body weight on the first day of testing that then stabilized but, importantly, there were no significant interactions of Day with Surgery, indicating that ORX and sham males similarly maintained body weight while on the PR1-CE task (Supplementary Figure S1).

**Figure 2.**
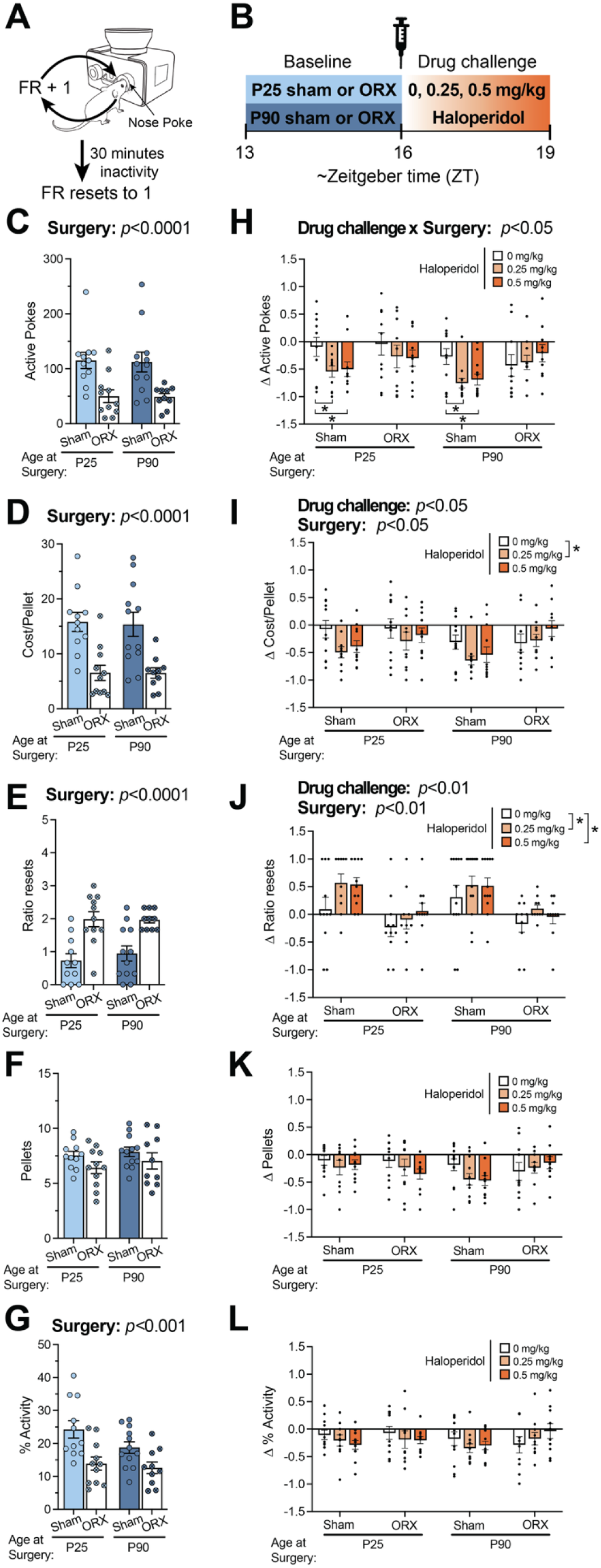
ORX increases effort cost sensitivity and attenuates behavioral effects of haloperidol. (**A**) Schematic of the PR1-CE task. The operant response requirement increased by 1 nose poke for each subsequent pellet and reset to 1 nose poke after 30 minutes of inactivity. (**B**) Timeline for drug challenge. Measures represent the mean hourly values during the 3-h baseline and 3-h post-injection periods, except for ratio resets **(E)**, which represent the total number of resets during each 3-h period. During baseline (**C-G**), sham mice (**C**) engaged in more active pokes, (**D**) paid a higher cost/pellet per hour (surgery: p’s<0.0001), and (**E**) exhibited fewer ratio resets (surgery: p<0.0001), while (**F**) earning a similar number of pellets compared to ORX mice. (**G**) Consistent with operant responding during baseline, shams engaged in more locomotor activity than ORX (surgery: p<0.001). In response to the drug challenge (**H-L**), haloperidol (**H**) reduced active pokes in shams (drug challenge x surgery: p<0.05), (**I**) reduced cost/pellet (drug challenge: p<0.05), and (**J**) increased ratio resets (drug challenge: p<0.01). Sham mice displayed a greater reduction in cost/pellet (surgery: p<0.05) and increase in ratio resets (surgery: p<0.01) than ORX mice. There were no significant changes in (**K**) pellets earned or (**L**) locomotor activity in response to the drug challenge. *p<0.05, Tukey’s

To examine dopaminergic contributions to PR1-CE performance, mice were administered vehicle or haloperidol (0.25 or 0.5 mg/kg; Figure 2B). Haloperidol reduced operant responding in sham mice but not ORX mice, reflected by a significant Surgery × Drug interaction for active pokes (F_2,77_=3.20, p=0.0461; Figure 2H). Post hoc analyses indicated that both doses reduced responding relative to vehicle in sham mice (p≤0.0003), whereas ORX mice showed minimal sensitivity (Figure 2H). Haloperidol also reduced cost/pellet (main effect of Drug: F_2,77_=3.64, p=0.0307; Figure 2I) and increased the frequency of ratio resets (F_2,77_=5.48, p=0.0059; Figure 2J), and sham mice exhibited greater changes in these measures compared to ORX mice (main effects of Surgery: F_1,40_=4.92, p=0.0323 and F_1,40_=17.67, p=0.0001, respectively). Pellets earned were unaffected (p>0.05; Figure 2K), and no group differences were observed in locomotor suppression (Figure 2L).

### Free Feeding and Sucrose Preference

To determine whether ORX- and haloperidol-induced changes in PR1-CE reflected altered effort sensitivity rather than nonspecific effects on consumption or reward palatability, we compared baseline and post-injection pellet consumption under free feeding conditions (Figure 3A,B). Sham mice took more pellets than ORX mice at baseline (main effect of Surgery: F_1,34_=6.43, p=0.0160), but haloperidol did not significantly alter pellets taken (p>0.05; Figure 3C,D). Baseline sucrose preference during SPT2 differed across groups (Surgery × Age interaction: F_1,41_=9.33, p=0.0040) but was unaffected by drug treatment (p>0.05; Figure 3E,F). Follow-up analyses revealed that P90 shams displayed a significantly lower sucrose preference compared to P90 ORX (F_1,41_=11.05, *p*=0.0019) and P25 sham counterparts (F_1,41_=9.38, *p*=0.0039) at baseline (Figure 3E). SPT1 and SPT3 results are provided in the supplemental materials. These findings indicate that ORX and haloperidol primarily affect effort-based responding rather than consummatory behavior or reward valuation.

**Figure 3.**
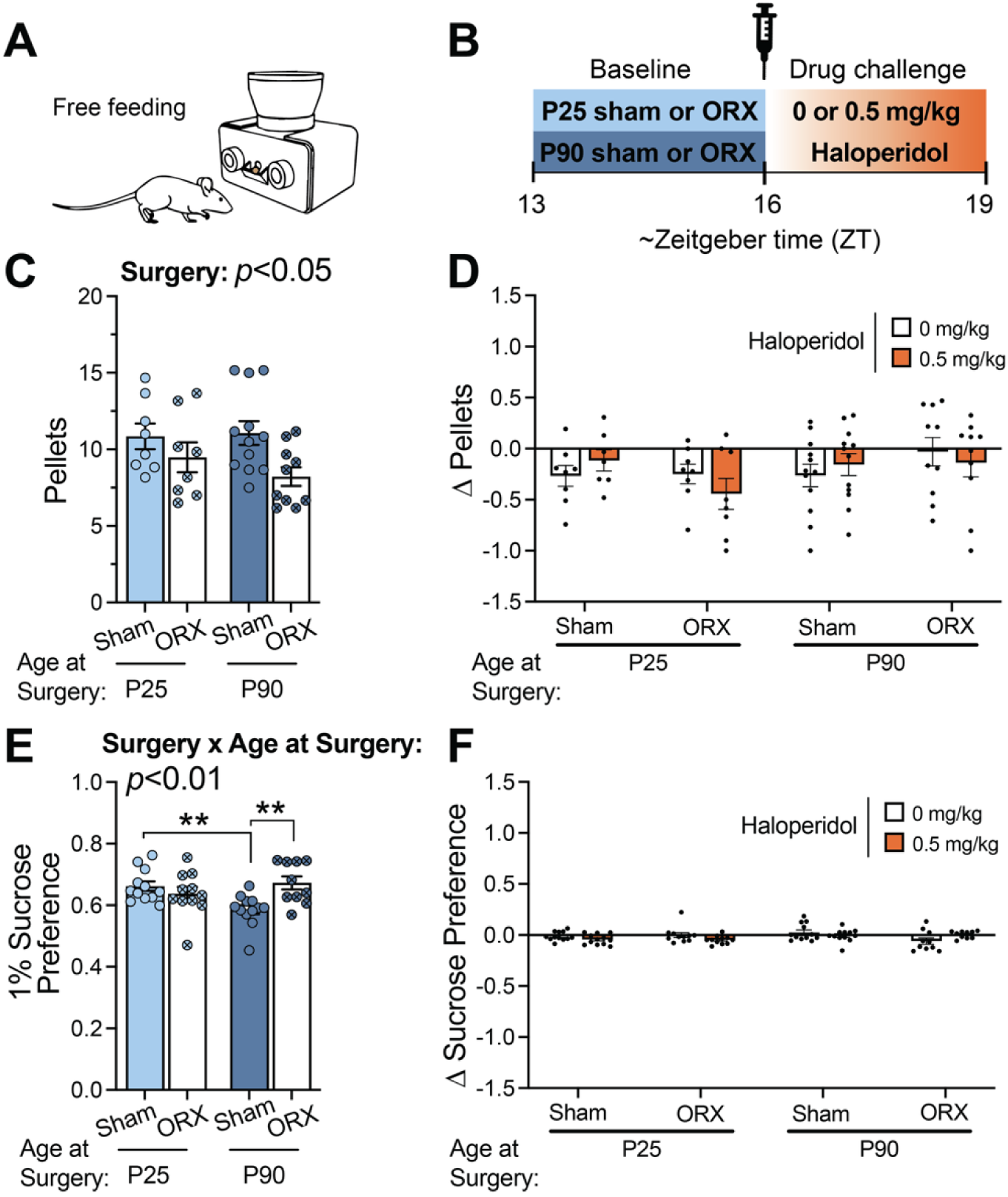
Haloperidol does not reduce pellet consumption or sucrose preference under no-cost conditions. (**A**) Schematic of the free feeding mode. A pellet was delivered each time one was removed from the magazine. (**B**) Timeline for drug challenge. At baseline, (**C**) sham mice took more pellets during free feeding than ORX mice (surgery: p<0.05), and (**D**) pellets taken were not impacted by haloperidol. During SPT2, (**E**) P90 shams displayed lower sucrose preference on the baseline day compared to P90 ORX and P25 shams, but (**F**) drug challenge did not alter sucrose preference. **p<0.01, simple main effects

### Dopamine Content

We next examined DA content in key striatal regions implicated in effort-based decision making, including dorsomedial striatum (DMS) and nucleus accumbens (NAc) (Figure 4A). DA content was higher in DMS than NAc overall (main effect of Region: F_1,52_=22.53, p<0.0001), but no effects of ORX were observed in the DMS (p>0.05; Figure 4B–E). In contrast, NAc DA content showed a significant Surgery × Age interaction (F_1,42_=4.26, p=0.0452), with postpubertal ORX mice exhibiting increased DA content relative to age-matched shams (p=0.0298) and prepubertal ORX mice (p=0.0360; Figure 4F). Metabolite analyses revealed corresponding reductions in NAc DOPAC:DA (F_1,39_=7.41, p=0.0096) and HVA:DA ratios (F_1,39_=4.94, p=0.0321) following postpubertal ORX (Figure 4G,H), consistent with decreased DA turnover. No effects were observed for 3-MT:DA ratios (p>0.05) (Figure 4I), and no significant changes were detected following prepubertal ORX.

**Figure 4.**
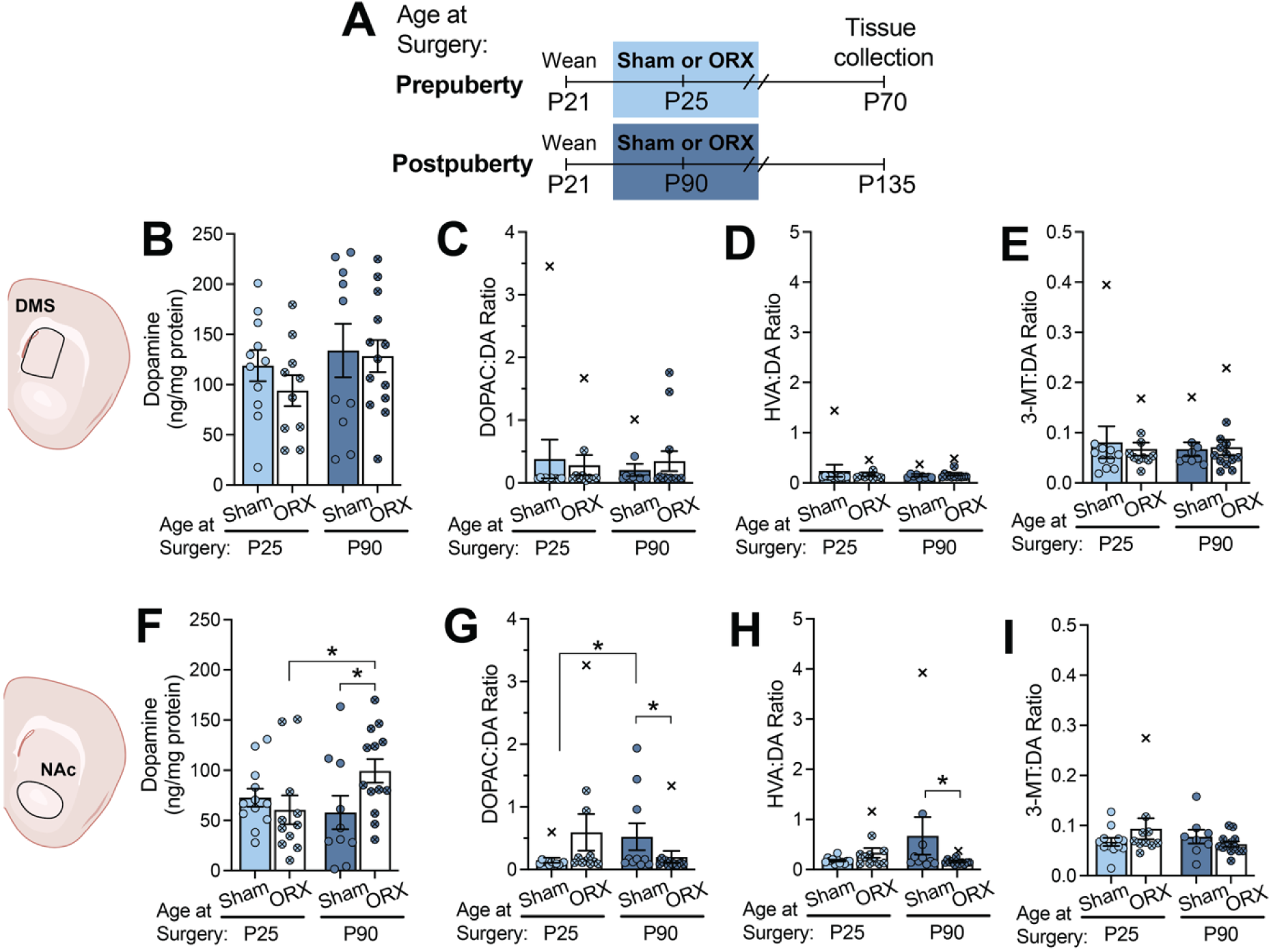
Post- but not prepubertal ORX alters nucleus accumbens dopamine measures. (**A**) Experimental timeline for Exp. 2 tissue collection for dopamine (DA) and metabolites levels. In the DMS (**B-E**), there were no group differences in dopamine levels or metabolite ratios. In the NAc (**F-I**), postpubertal ORX (**F**) increased DA content relative to age-matched shams and prepubertal ORX (age x surgery: p<0.05). The ratios of (**G**) DOPAC and (**H**) HVA to DA were reduced in postpubertal ORX relative to age-matched shams (age x surgery: p<0.01 and p<0.05, respectively). (**I**) 3-MT:DA was unaffected. Outliers (±5 standard deviations from the within group mean) were not included in statistical analyses and are shown as Xs. *p<0.05, simple main effects

Because ORX altered DA content in the NAc but not the DMS, we next examined whether these effects were associated with age-dependent changes in NAc circuit function. Using Drd1tdTomato reporter mice, we assessed the intrinsic excitability of DA D1 receptor-positive (D1R⁺) (Figure 5A–B) and D1R⁻ SPNs (Figure 6A–B) in the core region of the NAc following sham or ORX surgery performed at P25 or P90 using whole-cell patch-clamp recordings.

**Figure 5.**
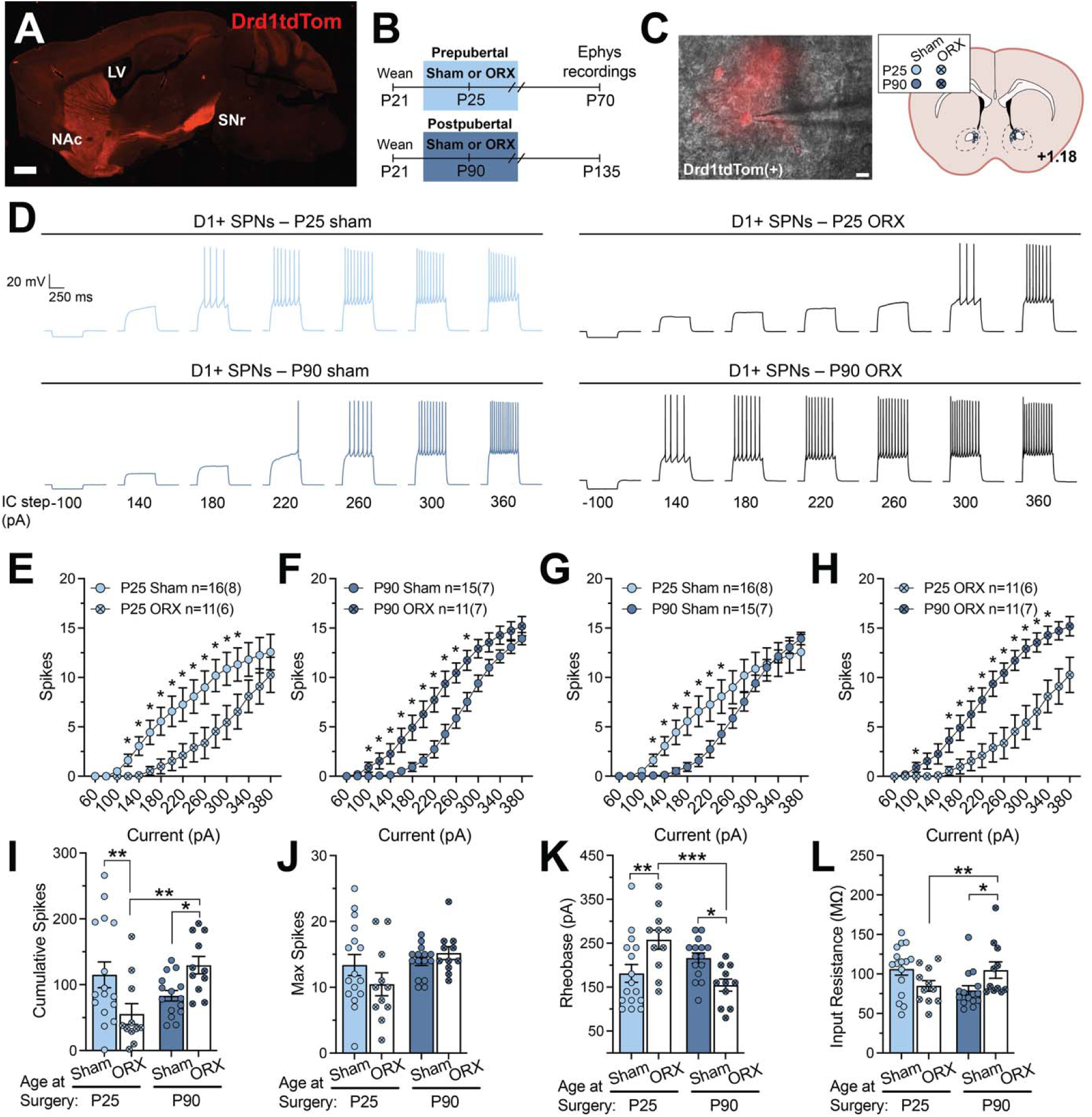
Pre- and postpubertal ORX differentially alter D1R^+^ SPN excitability in NAc core. (**A**) Sagittal overview of Drd1tdTomato expression in transgenic mouse. Scale bar represents 1 mm. (**B**) Experimental timeline for Exp. 3 electrophysiology. (**C**) Representative merged image of DIC and tdTomato showing pipette patched to D1R^+^ cell. Scale bar represents 10 μm. Atlas coronal slice representation of NAc core locations of recorded D1R^+^ cells by group. (**D**) Representative traces from current clamp recordings from D1R^+^ cells in the presence of synaptic blockers. ORX differentially impacted spike output of D1R^+^ cells across a range of currents depending on the timing (age x surgery x current: p<0.0001) (**E-H**). (**E**) Prepubertal ORX decreased spike output, while (**F**) postpubertal ORX increased spike output. (**G**) Older age was associated with reduced spike output among shams, while the opposite effect was seen among ORX mice (**H**). Other measures of intrinsic excitability (**I-L**) similarly indicated reduced excitability (**I**: reduced cumulative spikes, **K**: increased rheobase) after prepubertal ORX and enhanced excitability (**I**: increased cumulative spikes, **K**: decreased rheobase, and **L**: increased input resistance) after postpubertal ORX. (**J**) There were no group differences in maximum spikes. *p<0.05, **p<0.01, ***p<0.001, simple main effects

**Figure 6.**
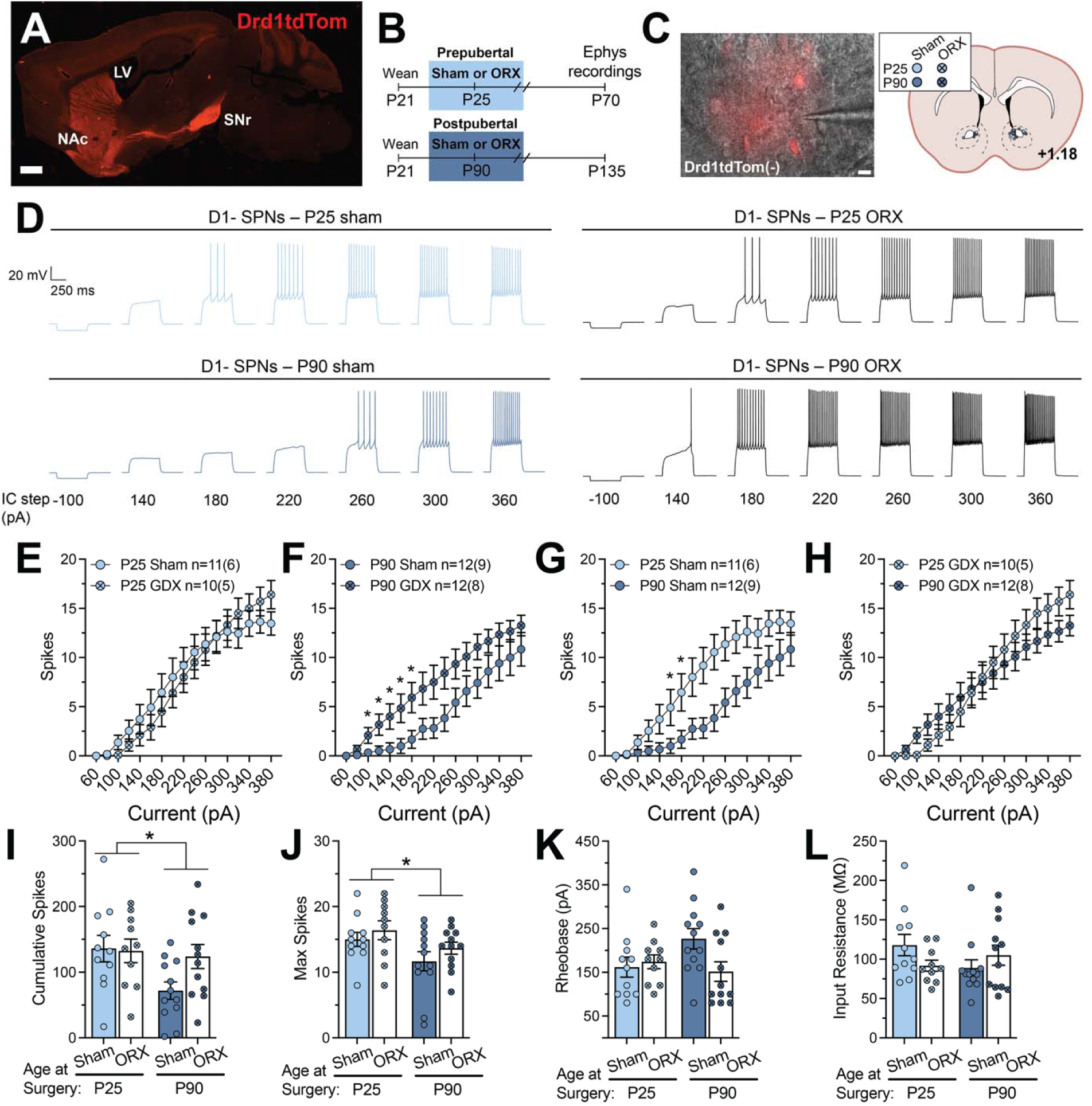
Pre- and postpubertal ORX differentially alter D1R^−^ SPN excitability in NAc core. (**A**) Sagittal overview of Drd1tdTomato expression in transgenic mouse. Scale bar represents 1 mm. (**B**) Experimental timeline for Exp. 3 electrophysiology. (**C**) Representative merged image of DIC and tdTomato showing pipette patched to D1R^−^ cell. Scale bar represents 10 μm. Atlas coronal slice representation of nucleus accumbens core locations of recorded D1R^−^ cells by group. (**D**) Representative traces from current clamp recordings from D1R^−^ cells in the presence of synaptic blockers. Timing of ORX significantly influenced its effects on D1R^−^ spike output at varying currents (age x surgery x current: p<0.001) (**E-H**). (**F**) Postpubertal ORX increased D1R^−^spike output, while (**E**) prepubertal ORX did not impact D1R^−^ spike output. (**G**) Older age was associated with reduced spike output among shams, while no age effects were seen among ORX mice (**H**). Cumulative spikes (**I**) and maximum spikes (**J**) were significantly reduced in older mice (age: *p’s<0.05), although this effect appears to be primarily driven by a reduction in P90 shams. Consistent with this, there was a trending surgery x age interaction (p=0.0521) for rheobase (**K**), reflecting increased rheobase in P90 shams compared to age-matched ORX mice (p<0.05) and P25 shams (p<0.05). (**L**) There were no significant group differences in input resistance. *p<0.05, simple main effects

### Dopamine D1R^+^ Cells

Intrinsic excitability of D1R^+^ SPNs was differentially affected by ORX depending on developmental timing (age x surgery x current: p<0.0001; Figure 5E–H). Prepubertal ORX reduced excitability, as indicated by fewer cumulative evoked spikes (age x surgery: F_1,49_=7.47, *p*=0.0087; Figure 5I) but no effect on max spikes (Figure 5J), increased rheobase (age x surgery: F_1,49_=14.95, p=0.0003; Figure 5K), and reduced input resistance (age x surgery: F_1,49_=8.78, p=0.0047; Figure 5L). In contrast, postpubertal ORX increased excitability relative to P90 shams, reflected by greater spike output, reduced rheobase, and increased input resistance (p’s<0.05). No differences were observed in resting membrane potential (Supplementary Figure S2A).

### Dopamine D1R^−^ Cells

Intrinsic excitability of D1R^−^ SPNs was also differentially affected by ORX depending on developmental timing, but showed a different pattern compared to D1R^+^ SPNs (Figure 6E–H). Prepubertal ORX did not significantly alter excitability (p>0.05) (Figure 6E), whereas postpubertal ORX slightly increased excitability (Figure 6F). A shift towards greater D1R^−^ SPN excitability in P90 ORX mice was also reflected by a trending surgery x age interaction (F_1,41_=4.00, *p*=0.0521) for rheobase with simple main effect analyses indicating reduced rheobase in P90 ORX compared to P90 shams (F_1,41_=6.35, *p*=0.0157; Figure 6K). Moreover, older age was associated with increased rheobase among shams (F_1,41_=4.54, *p*=0.0391; Figure 6K). A main effect of age was observed for cumulative spikes (F_1,41_=4.31, *p*=0.0442; Figure 6I) and max spikes (F_1,41_=5.98, *p*=0.0189; Figure 6J), with greater excitability in P25 mice overall. Taken together, it appears that this age-related decrease in excitability was primarily driven by a reduction in the P90 sham group, as spike output from P90 ORX was more in line with that of the prepubertal groups. No differences were observed in resting membrane potential (Supplementary Figure S2B).

## Discussion

The current study investigated how testicular hormones modulate mesoaccumbal circuits to regulate reward pursuit and effort sensitivity. Consistent with prior work showing that high-dose testosterone reduces sensitivity to increasing effort requirements (Wallin et al., 2015), we found that androgen depletion caused by ORX had the opposite effect, markedly increasing effort sensitivity and biasing animals towards an energy-efficient behavioral strategy. In the PR1-CE task, ORX mice maintained overall food intake despite more frequent ratio resets and a lower cost/pellet, indicating preserved reward acquisition alongside reduced willingness to sustain high-effort responding. This pattern parallels established effects of dopaminergic signaling, whereby reduced DA function increases effort sensitivity (Beeler et al., 2012b, 2012c), suggesting that androgen loss shifts mesoaccumbal DA function toward a lower-gain, energy-conserving state.

After establishing baseline differences in PR1-CE performance, we probed the contribution of D2R signaling, which has been shown to regulate effort sensitivity (Mourra et al., 2020). Systemic administration of haloperidol, a dopamine antagonist that primarily acts at D2R, reduced effort expenditure in sham mice but had minimal effects in ORX mice, indicating reduced behavioral sensitivity to dopaminergic perturbation following androgen depletion. This finding complements prior work showing that testosterone enhances sensitivity to D2R antagonism (Donovan and Wood, 2022) and that testosterone’s ability to rescue voluntary wheel-running in ORX mice is dependent on D2R signaling (Jardí et al., 2018), suggesting that androgens bidirectionally modulate dopaminergic control of effortful behavior. Together, these data support the interpretation that androgen loss diminishes the functional impact of dopamine signaling during effortful reward pursuit.

Our behavioral and pharmacological findings are supported by neurochemical changes within the NAc. Postpubertal ORX increased DA content while reducing DOPAC:DA and HVA:DA ratios, a pattern consistent with reduced DA turnover (Lookingland and Moore, 2005). Prior work demonstrates that testicular hormones regulate multiple aspects of striatal DA signaling, including synthesis, release, and reuptake (Abreu et al., 1988; Jackson et al., 2025; Shemisa et al., 2006). Our observed neurochemical changes are consistent with the attenuated behavioral response to D2R antagonism and increased effort sensitivity observed in ORX mice.

Recent work provides a circuit-level framework for interpreting the effects of ORX on NAc SPNs. Walle et al. demonstrated that the balance of activity between NAc core D1R^+^ vs. D2R^+^ SPN populations regulates energy allocation towards reward pursuit and physical activity (Walle et al., 2024). More specifically, D1R^+^ SPN activity facilitated effortful responding for food and voluntary exercise (Walle et al., 2024). These findings are in line with another recent study that showed that obese mice – who expend more physical effort to earn food – exhibit enhanced intrinsic excitability and task-related *in vivo* activity of D1R+ SPNs in the NAc core (Matikainen-Ankney et al., 2023). Consistent with this framework, prepubertal ORX selectively reduced D1R+ SPN excitability, providing a plausible mechanism for the observed reduction in effortful responding despite preserved food intake. In contrast, postpubertal ORX increased the excitability of both D1R^+^ and D1R^−^ (putative D2R^+^) SPNs, suggesting a more complex reconfiguration of NAc circuit function. While this pattern does not indicate a simple reduction in D1R^+^/D2R^+^ SPN balance, it may suggest a change in how these populations are recruited during behavior. For instance, the reduced dopamine turnover observed in postpubertal ORX mice may limit phasic dopamine signaling that is required to effectively engage lower-affinity D1R and compress the dynamic range of dopamine “seen” by D2R, potentially contributing to reduced behavioral sensitivity to D2R antagonism (Dreyer et al., 2010). Together with our behavioral and neurochemical findings, these results suggest that androgen loss shifts the functional balance of NAc output toward a thriftier mode of energy allocation, with developmental timing determining whether this occurs via selective D1R^+^ SPN hypofunction or broader alterations in network excitability. Future studies examining *in vivo* dynamics of D1R^+^ and D2R^+^ population activity within NAc will be critical for linking these intrinsic changes caused by ORX to circuit activity during food-seeking and locomotor behaviors.

Emerging evidence indicates that pubertal hormone exposure can shape the long-term function of neural systems involved in affective and motivational processing (Boivin et al., 2017; Delevich et al., 2020a; Jackson et al., 2025; Klappenbach et al., 2023; Schulz and Sisk, 2016; Wright et al., 2023). Although we observed similar behavioral outcomes following ORX before or after puberty, these effects were associated with distinct neurochemical and cellular adaptations. Postpubertal ORX was associated with increased dopamine content and reduced metabolite-to-dopamine ratios in the NAc, whereas prepubertal ORX selectively reduced D1R^+^ SPN excitability without altering dopamine levels. Together, these findings suggest that androgen loss produces a common behavioral outcome – enhanced effort cost sensitivity – via distinct mechanisms depending on developmental timing. Future studies incorporating hormone replacement during puberty vs. later in adulthood are required to determine whether the observed developmental timing of effects are consistent with organizational effects of pubertal hormones on NAc circuitry that differ from activational modulation in adulthood.

### Limitations and Future Directions

Our study has a number of limitations. Cost sensitivity encompasses multiple domains beyond physical effort, including delay, uncertainty, punishment, and cognitive effort. Although testosterone has been shown to modulate several of these domains in a domain-specific manner, our study focused exclusively on effort-based costs, and cannot determine whether androgen loss similarly influences other forms of cost-benefit decision making (Dokovna et al., 2019; Wallin et al., 2015; Wood et al., 2024, 2013). Because increasing response requirements in the PR1-CE task also prolong reward delay, we also cannot fully dissociate effort from temporal cost sensitivity.

We did not perform hormone replacement experiments to determine whether testosterone or its metabolite estradiol are sufficient to reverse the observed behavioral and neural effects. In addition, ORX does not eliminate locally synthesized neurosteroids, and residual AR/ER signaling may have influenced our results (Seib et al., 2023; Tobiansky et al., 2018b). While we observed alterations in NAc dopamine content and SPN intrinsic excitability, we did not directly assess dopamine receptor signaling or its contribution to these cellular changes. Furthermore, our HPLC measurements lacked the spatial resolution to distinguish NAc subregions, whereas electrophysiological recordings were restricted to the core. Finally, our neurochemical findings are consistent with altered dopamine turnover, but do not capture the dynamics of dopamine release and reuptake during behavior. Future studies using *in vivo* dopamine sensors will be necessary to link androgen loss to real-time dopamine signaling and motivated behavior.

## Conclusion

Androgen depletion caused by ORX increases sensitivity to effort-related costs, biasing behavior toward energy-efficient strategies while preserving food intake. This phenotype is consistent with a shift in mesoaccumbal dopamine function and is supported by convergent pharmacological, neurochemical, and electrophysiological evidence. Notably, similar behavioral outcomes following pre- and postpubertal ORX were mediated by distinct circuit-level adaptations, indicating that developmental timing shapes the neural implementation rather than the expression of effort-based decision making. These findings support a role for testicular hormones in regulating mesoaccumbal circuits involved in effort-based decision making and provide a framework for understanding motivational deficits associated with male hypogonadism.

## Acknowledgments

The authors would like to thank Tyler Edwards for skilled technical support during the behavioral study. Additionally, the authors would like to thank Vanderbilt University’s Neurochemistry Core Laboratory for their excellent technical assistance with analyzing tissue samples for neurochemical content. Finally, we would like to thank Dr. Lex Kravitz and FedForum members for technical assistance. This study was supported by a Young Investigator Grant from the Brain & Behavior Research Foundation and an Intramural Grant from the College of Veterinary Medicine at Washington State University (to KMD). SRW was supported by an NIH F32 fellowship from the National Institute on Drug Abuse (F32DA060685) and the State of Washington Initiative Measure No. 502 (SRW, KMD, RJM).

