## Supplementary Materials for "Androgen depletion increases sensitivity to effort-related costs and alters mesoaccumbal circuit function in male mice"

Westbrook *et al.*

**Supplemental Methods**

Animals and Surgery

The current study used 95 male C57BL/6 mice (Experiment 1: n=48; Experiment 2: n=47) and 31 male B6.Cg-Tg(Drd1a-tdTomato)6Calak/J mice (Experiment 3; JAX stock #016204 (Ade et al., 2011)) generated in-house from breeders originally obtained from Charles River and Jackson Laboratory, respectively. Mice were weaned on postnatal day (P) 21, group housed and maintained on a reverse 12-hour light/dark cycle (lights off at 0700). Mice underwent prepubertal (P25) or postpubertal (P90) orchiectomy (ORX) or sham surgeries as previously described (Klappenbach et al., 2023). For Experiment 1, mice were individually housed beginning 35 days after surgery. Experiments 2 and 3 used group-housed mice with ad libitum access to chow (Purina 5001) and water. All procedures were approved by the Institutional Animal Care and Use Committee at Washington State University and followed AAALAC and NIH guidelines.

Exp. 1: Behavioral Procedures

*Food Intake and Operant Responding*

Feeding Experimentation Devices (FED3; Matikainen-Ankney et al., 2021) continuously recorded operant responding and food intake (20 mg grain-based pellets; Test Diet 5-TUM) in the home cage and served as the sole source of food beginning 35 days after surgery. During FED3 testing, mice were individually housed in a cage with an isopad (Braintree Scientific, Inc) to prevent obstruction of pellet dispensers. Mice and FED3 dispensers were weighed daily and the number of pellets dispensed was recorded.

FED3 dispensers were initially set to free-feeding mode, in which a pellet was dispensed whenever a pellet was removed from the magazine. After at least three days in free-feeding mode, FED3 dispensers were switched to fixed-ratio 1 (FR1), during which one poke to the active port (left) earned one pellet. After at least 2 days on FR1, mice were switched to a progressive ratio 1 closed economy (PR1-CE) schedule. Under PR1-CE conditions, the response requirement increased by one active nose poke for each subsequent pellet and reset to 1 nose poke following 30 minutes of inactivity (ratio reset). Mice remained on PR1-CE for the majority of the study. After a series of sucrose preference testing and drug challenges described below, FED3 dispensers were returned to free-feeding mode before the final round of sucrose preference testing and drug challenge.

*Sucrose Preference Testing (SPT)*

Baseline sucrose preference testing (SPT1) was conducted after at least three days of PR1-CE. For the first two days, mice were provided with two water bottles to habituate to the testing procedure and bottle weights were recorded every 24 hours to measure consumption. During the subsequent two days, one bottle contained water and the other 1% (w/v) sucrose. Bottle locations were switched after 24 hours.

*Locomotor Activity*

Home-cage locomotor activity was continuously monitored using Pallidus MR1 passive infrared (PIR) activity sensors, which detect movement of warm objects within the home cage (Usiyevich et al., 2025). Activity data were collected and binned in 1-h intervals, and PIR activity sensors were placed in the home cages at least 24 hours before the first drug challenge session.

*Drug Challenge*

Haloperidol (Sigma-Aldrich H1512) was dissolved in 0.3% tartaric acid (Yun et al., 2023) to a final concentration of 0, 0.025, or 0.05 mg/ml for final injection volumes of 10 ml/kg across doses (0, 0.25, or 0.5 mg/kg). The first drug challenge period occurred during PR1-CE after baseline sucrose preference testing. Sucrose was not present during this period. Mice received 0, 0.25, and 0.5 mg/kg haloperidol injections (i.p.) at 1100 (ZT16 ± 1.5) in ascending order across days with at least 48 hours between injections. For the second and third drug challenges, only 0 and 0.5 mg/kg doses were used, and dose order was counterbalanced. The second drug challenge period occurred during PR1-CE, and the third and final drug challenge occurred during FF. The second and third challenges occurred during periods with simultaneous sucrose preference testing (SPT2 and SPT3), and injections occurred on the second day of sucrose preference testing.

*Seminal Vesicle Collection*

At least one day after the completion of behavioral testing, mice were euthanized via CO_2_. Cardiac punctures were performed for blood collection and serum was stored at -80°C. Seminal vesicles were dissected and weighed to verify that ORX reduced circulating testosterone levels.

*Behavioral Data Analysis*

*Food Intake and Operant Responding.* FED3 data were written to a secure digital (SD) card on each behavioral device, and custom R scripts were used to quantify active and inactive pokes, pellets taken, and ratio resets. All PR1-CE data from one mouse (P25 sham) was excluded due to multiple instances of FED equipment error. PR1-CE and locomotor activity data from two mice were excluded for the 0.25 mg/kg haloperidol injection day due to FED equipment error (n=2, P25 ORX). Data from one mouse (P25 sham) for the 0.25 mg/kg haloperidol injection day was more than 5 standard deviations from the mean of the group and was excluded from analyses. Data from the remaining 44 mice were analyzed (n’s=10-12/group).

The first cohort of mice (n=4 P25 sham; n=3 P25 ORX) did not undergo the final phase of the experiment (i.e., free-feeding sessions with drug challenge and SPT3). For one mouse (P25 sham) free-feeding data after 0.5 mg/kg haloperidol injection was more than 5 standard deviations from the group mean, so data from this mouse were excluded from analyses. Data from the remaining 38 mice were analyzed (n’s=7-12/group).

*Sucrose Preference Testing.* For SPT1, data was excluded for 2 mice (n=1 P25 sham, n=1 P25 ORX) due to equipment error. For SPT2, there were no outliers or equipment errors, so data from all 45 mice were included in sucrose preference analyses. As noted above, the first cohort of mice (n=4 P25 sham; n=3 P25 ORX) did not undergo SPT3. Sucrose preference was calculated as sucrose consumed/(sucrose + water consumed).

*Baseline*. During the 3-h period preceding each drug challenge, cost/pellet per hour, active pokes per hour, pellets earned per hour, ratio resets, and activity per hour were analyzed using 3-way (Day x Surgery x Age at Surgery) mixed ANOVAs, with Day as the within-subjects factor. For sucrose preference, the 24-h period preceding each drug challenge was used as the baseline measure. Because there were no significant main effects or interactions involving Day during baseline, data were collapsed across Day and reanalyzed using 2-way (Surgery x Age at Surgery) ANOVAs. Baseline values presented in Figure 2C-G represent averages per animal collapsed across Day. Significant interactions were followed by simple main effects analyses.

*Drug Challenge.* For each drug challenge (0, 0.25, or 0.5 mg/kg haloperidol), drug-induced changes in cost/pellet, active pokes, pellets earned, ratio resets, and activity were quantified using a normalized change score calculated as (post-injection – pre-injection)/(post-injection + pre-injection). Pre- and post-injection values were derived from the 3-h periods immediately before and after injection, respectively. Cost/pellet, active pokes, pellets earned, and activity were calculated as hourly values and averaged across each 3-h period, whereas ratio resets were summed across each 3-h period. For sucrose preference, change scores were calculated using 24-h pre- and post-injection measurements. Change scores were analyzed using 3-way (Drug Challenge x Surgery x Age at Surgery) mixed ANOVAs with Drug Challenge as the within-subjects factor. Significant interactions were followed by simple main effects analyses.

Exp. 2: Dopamine Content

C57BL/6 mice that underwent prepubertal (P25) surgery were euthanized on P70 and those that underwent postpubertal (P90) surgery were euthanized on P135. Mice were rapidly decapitated, and brains were placed in a matrix and sectioned into 1 mm coronal slices. Sections were maintained on dry ice during tissue collection, and 1 mm^3^ punches were collected from the nucleus accumbens (NAc) and dorsomedial striatum (DMS) using anatomical landmarks corresponding to approximately Bregma +1.77 to +0.73 mm and +0.95 to +0.5 mm, respectively. Punches from both hemispheres were pooled within each mouse, snap frozen in liquid nitrogen, and stored at -80°C until shipment to Vanderbilt University’s Neurochemistry Core Laboratory. Dopamine and its metabolites (3,4-dihydroxyphenylacetic acid [DOPAC], homovanillic acid [HVA], and 3-methoxytyramine [3-MT]) were quantified using high-performance liquid chromatography with electrochemical detection.

*Dopamine Data Analysis*

Dopamine, DOPAC, HVA, and 3-MT content were measured as ng/mg protein, and the ratio of each metabolite (DOPAC, HVA, and 3-MT) to dopamine was calculated. There were no outliers for dopamine content. Outliers (± 5 standard deviations from the group mean) for dopamine metabolites were excluded from analyses and are shown in Figure 4 as X’s. Two DMS samples were lost in transit (n=1 P25 ORX, n=1 P25 sham). Measures where the value was below the limit of detection for a sample were excluded from statistical analyses. The final sample sizes per measure are provided in Table S1. Three-way (Surgery, Age at Surgery, Brain Region) mixed ANOVA was conducted on dopamine content with brain region (DMS, NAc) as the within-subject factor. There was a significant main effect of brain region, so further analyses (2-way Surgery x Age at Surgery ANOVAs) on dopamine content and metabolite ratios were conducted separately for each brain region. Where significant interactions were detected, simple main effects analyses were performed.

Exp. 3: Electrophysiology

*Slice preparation.* B6.Cg-Tg(Drd1a-tdTomato)6Calak/J mice were deeply anesthetized with an overdose of ketamine/xylazine and transcardially perfused with ice-cold cutting buffer containing (in mM): 110 choline-Cl, 2.5 KCl, 7 MgCl_2_, 0.5 CaCl_2_, 25 NaHCO_3_, 11.6 Na-ascorbate, 3 Na-pyruvate, 1.25 NaH_2_PO_4_, and 25 D-glucose, and bubbled in 95% O_2_/5% CO_2_. Coronal sections (300 µm) were prepared using a vibratome and transferred to oxygenated aCSF containing (in mM): 120 NaCl, 2.5 KCl, 1.3 MgCl_2_, 2.5 CaCl_2_, 26.2 NaHCO_3_, 1.25 NaH_2_PO_4_ and 11 D-glucose. Slices were bubbled with 95% O_2_/5% CO_2_ and incubated for 30 min in a 37°C bath. Following 30 min of recovery at room temperature, slices were transferred to the recording chamber. Recordings were made using a Multiclamp 700B amplifier (Molecular Devices), and analog signals were low-pass filtered at 2 kHz and digitized at 10 kHz with a Digidata 1550B interface (Molecular Devices). Clampex software (v11) was used for data acquisition. Only cells with an access resistance <25 MΩ were included in analyses. Access resistance and liquid junction potentials were not corrected. Whole-cell current clamp recordings were obtained from Drd1tdTomato+ and Drd1tdTomato- cells in the NAc core. Borosilicate electrodes (2.5-4.5 MΩ) contained potassium gluconate internal solution consisting of (in mM): 145 K Gluconate, 3 KCl, 10 HEPES, 0.2 EGTA, 2 MgCl_2_, and 2 Na_2_ATP. Recordings were performed at native resting membrane potential in the presence of NBQX (10 µM; HelloBio HB0443), DL-AP5 (10 μM; HelloBio HB0252), and GABAzine (10 μM; HelloBio HB0901). Neurons were injected with 500-ms current steps starting at -100 pA and increasing by 20 pA steps until a maximum of 380 pA. A 100 pA hyperpolarizing current step (30 sweeps) was applied to measure the input resistance based on the steady-state response.

*Electrophysiology Data Analysis*

Dependent measures from the current clamp recordings included spikes, cumulative spikes, maximum spikes, rheobase (pA), input resistance (MΩ), and resting membrane potential. Dependent measures from D1R^+^ and D1R^-^ cells were analyzed separately, except where noted below for depolarization block. Spikes were analyzed using 3-way (Surgery x Age at Surgery x Current) mixed ANOVA with Current as the within-subjects factor. Two-way (Surgery x Age at Surgery) ANOVAs were conducted for each of the other dependent measures. Where significant interactions were detected, simple main effects analyses were performed.

**Supplementary Results**

Exp. 1: Behavior

*Sucrose Preference Testing*

SPT1 occurred simultaneously with PR1-CE, but prior to any drug challenge in this experiment; thus, there was only a baseline measure. For SPT1, the effect of surgery depended on age (surgery x age interaction: F_1,39_=6.14, *p*=0.0176; Figure S3A). ORX significantly reduced sucrose preference relative to shams when surgery occurred before, but not after, puberty (simple main effect of surgery within P25 group: F_1,39_=6.20, *p*=0.0171). By SPT2, the prepubertal ORX group increased their sucrose preference to a comparable level to their sham counterparts. This change in sucrose preference in the prepubertal ORX group could be indicative of a failure to detect sucrose or a transient anhedonia that resolved by SPT2 (~10 days later). SPT1 was the only test that first began with two water bottles intended to get the mice accustomed to having access to two bottles; however, this practice may have differentially impacted our groups. We have previously reported less task engagement in a foraging task that requires exploration and approach in prepubertally ORX male mice (Delevich et al., 2020). Thus, it is possible that reduced exploration and or novelty aversion in this group may have resulted in a lack of or a later discovery of the change to sucrose from water in one of the bottles. An alternative explanation could be a transient anhedonia after ORX; however, it is unclear why we did not see this anhedonia in the postpubertal ORX group as this ORX timing would be in line with other reports (Carrier et al., 2015; Carrier and Kabbaj, 2012). Another explanation could be a delayed maturation of heightened reward sensitivity in young adulthood in prepubertal ORX mice. Rodent studies have shown a peak in reward sensitivity in adolescence and the peripubertal period (Friemel, 2010; Wilmouth and Spear, 2009); however, our mice at SPT1 are ~P70, which may be considered young adulthood but likely not adolescence.

SPT3 occurred simultaneously with FF and drug challenge, so it was analyzed at baseline and as a change score. For SPT3 baseline, 3-way ANOVA including Day failed to indicate any significant main effects or interactions with Day, so 2-way (surgery x age) ANOVA was conducted collapsed across Day. Younger mice had significantly higher sucrose preference compared to older mice [main effect of age: F_1,34_=6.61, *p*=0.0147; Figure S3B]. There was no significant main effect or interaction with surgery. Three-way (drug x surgery x age) ANOVA failed to reveal any significant main effects or interactions for change score during the drug challenge (Figure S3C), indicating that haloperidol challenge did not impact sucrose preference.

Exp. 3: Electrophysiology

Depolarization block was defined as a decrease in spiking with increasing current injections. Some cells underwent depolarization block as shown in Table S2. A 2 (Age at Surgery) x 2 (Surgery) x 2 (Cell Type) x 2 (Depolarization Block) Fisher’s exact test indicated no significant effect of cell type on the likelihood of undergoing depolarization block within any of the groups, so cell type was collapsed. A 2 (Age at Surgery) x 2 (Surgery) x 2 (Depolarization Block) Fisher’s exact test revealed significant associations. Specifically, significantly more cells from P25 shams underwent depolarization block compared to P25 ORX mice [*p*=0.0121] and P90 sham mice [*p*=0.0252]. No other comparisons were statistically significant.

**Supplementary Tables**

**Supplementary Table S1**. Final sample sizes per group for each measure of neurochemical content in the NAc and DMS.

|  |  | **P25 Sham** | **P25 ORX** | **P90 Sham** | **P90 ORX** |
| --- | --- | --- | --- | --- | --- |
| **Nucleus Accumbens** | **Dopamine** | 12 | 11 | 10 | 13 |
|  | **DOPAC:DA** | 11 | 10 | 10 | 12 |
|  | **HVA:DA** | 12 | 10 | 9 | 12 |
|  | **3-MT:DA** | 12 | 10 | 8 | 13 |
| **Dorsomedial Striatum** | **Dopamine** | 11 | 10 | 10 | 13 |
|  | **DOPAC:DA** | 10 | 9 | 9 | 13 |
|  | **HVA:DA** | 10 | 8 | 9 | 12 |
|  | **3-MT:DA** | 10 | 9 | 8 | 12 |

**Supplementary Table S2**. Sample sizes per group of D1R^+^ and D1R^-^ cells in the nucleus accumbens core that did or did not undergo depolarization block.

|  | **Depolarization Block?** | **P25 Sham** | **P25 ORX** | **P90 Sham** | **P90 ORX** |
| --- | --- | --- | --- | --- | --- |
| **D1R^+^** | **Yes** | 4 | 0 | 0 | 0 |
|  | **No** | 12 | 11 | 15 | 11 |
| **D1R^-^** | **Yes** | 3 | 0 | 1 | 1 |
|  | **No** | 8 | 10 | 11 | 11 |

**Supplementary Figures**

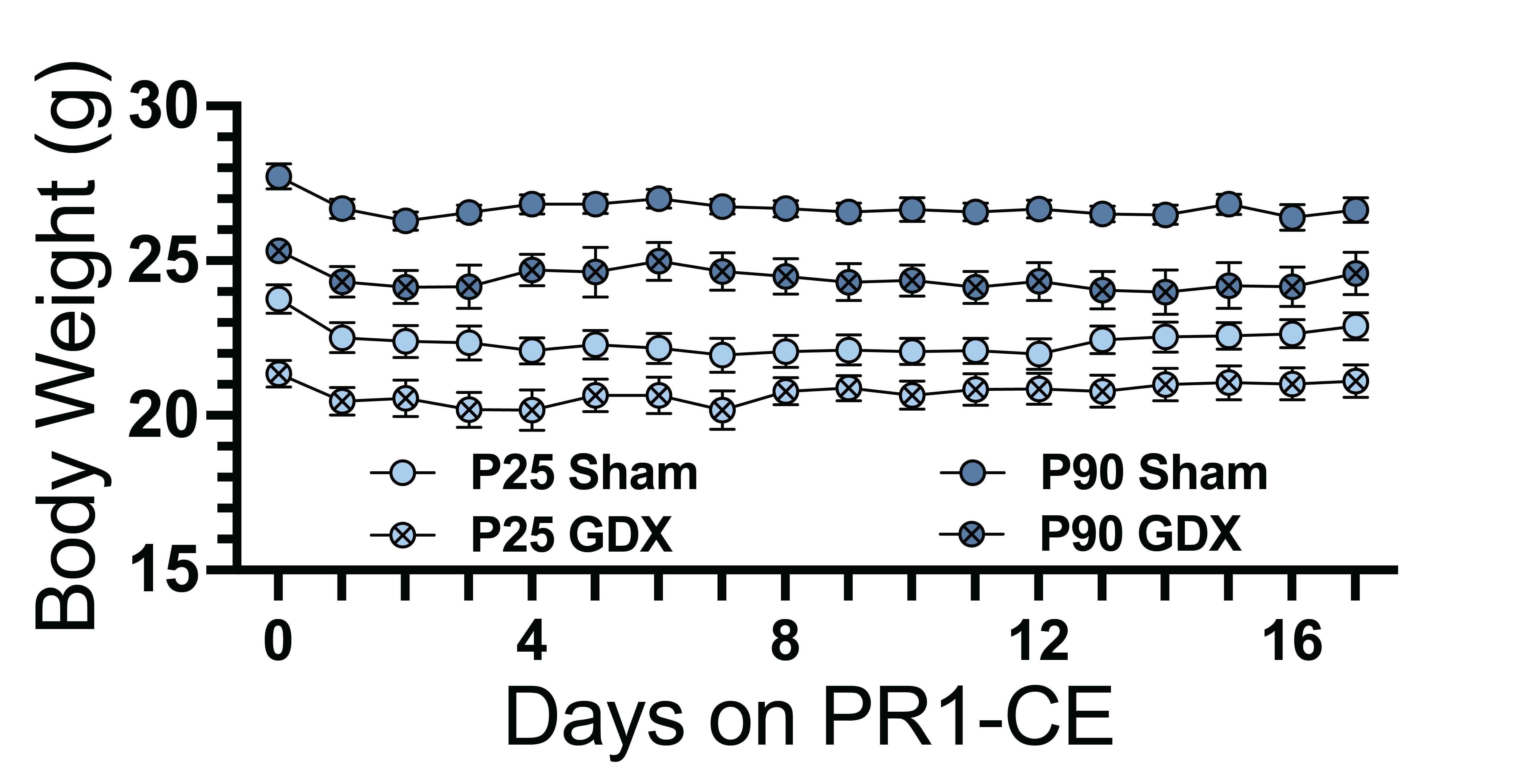

**Supplementary Figure S1**. Body weight (g) across 17 days on PR1-CE. Shams weighed more than ORX mice (surgery: p<0.0001), and older mice weighed more than younger mice (age: p<0.0001). Body weights significantly dropped for all groups on day 1 (day: p<0.0001), but these group differences were maintained throughout this period.

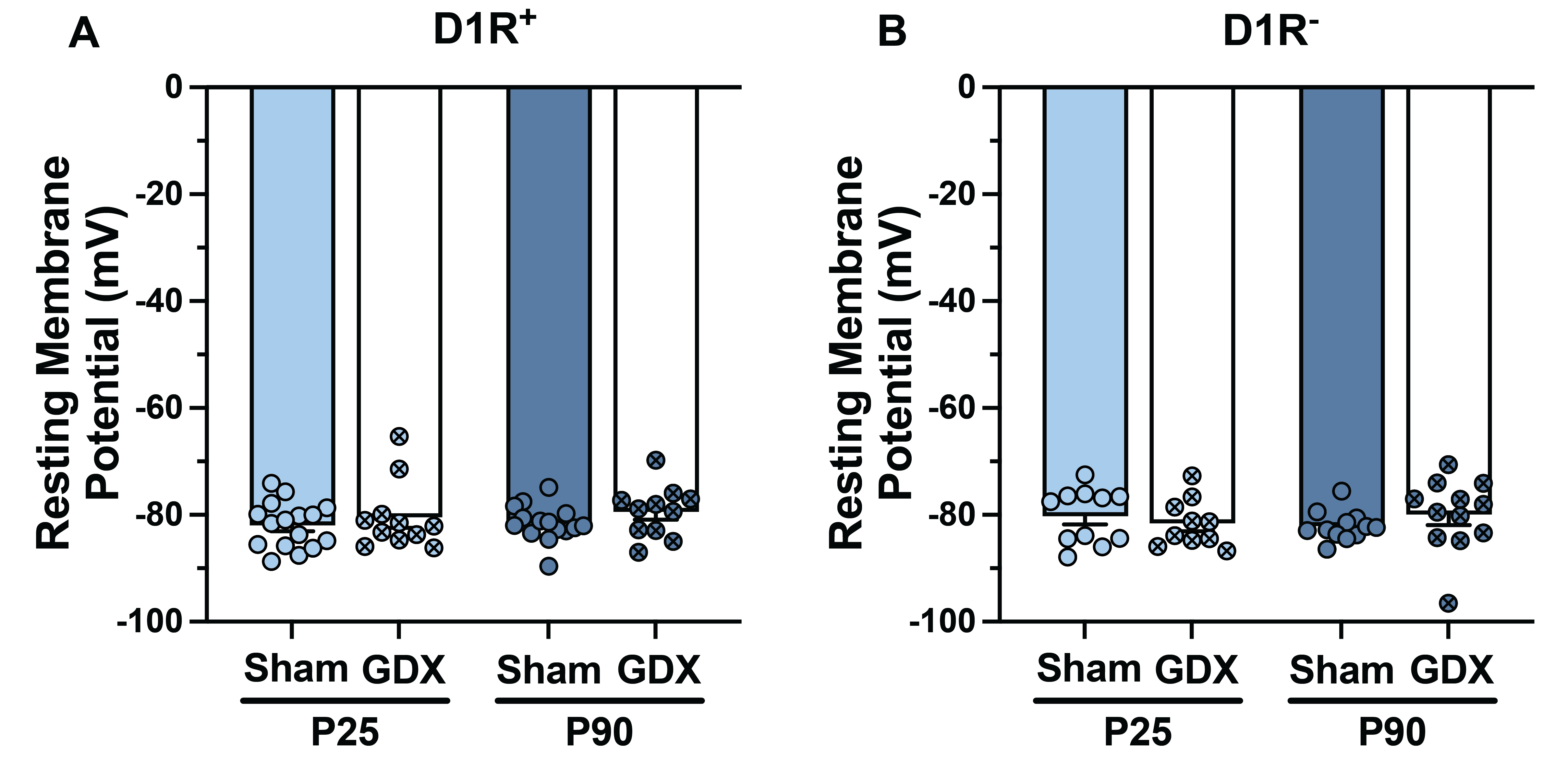

**Supplementary Figure S2**. Resting membrane potential (mV) of (**A**) D1R^+^ and (**B**) D1R^-^ SPNs. There were no group differences in resting membrane potential for either cell type.

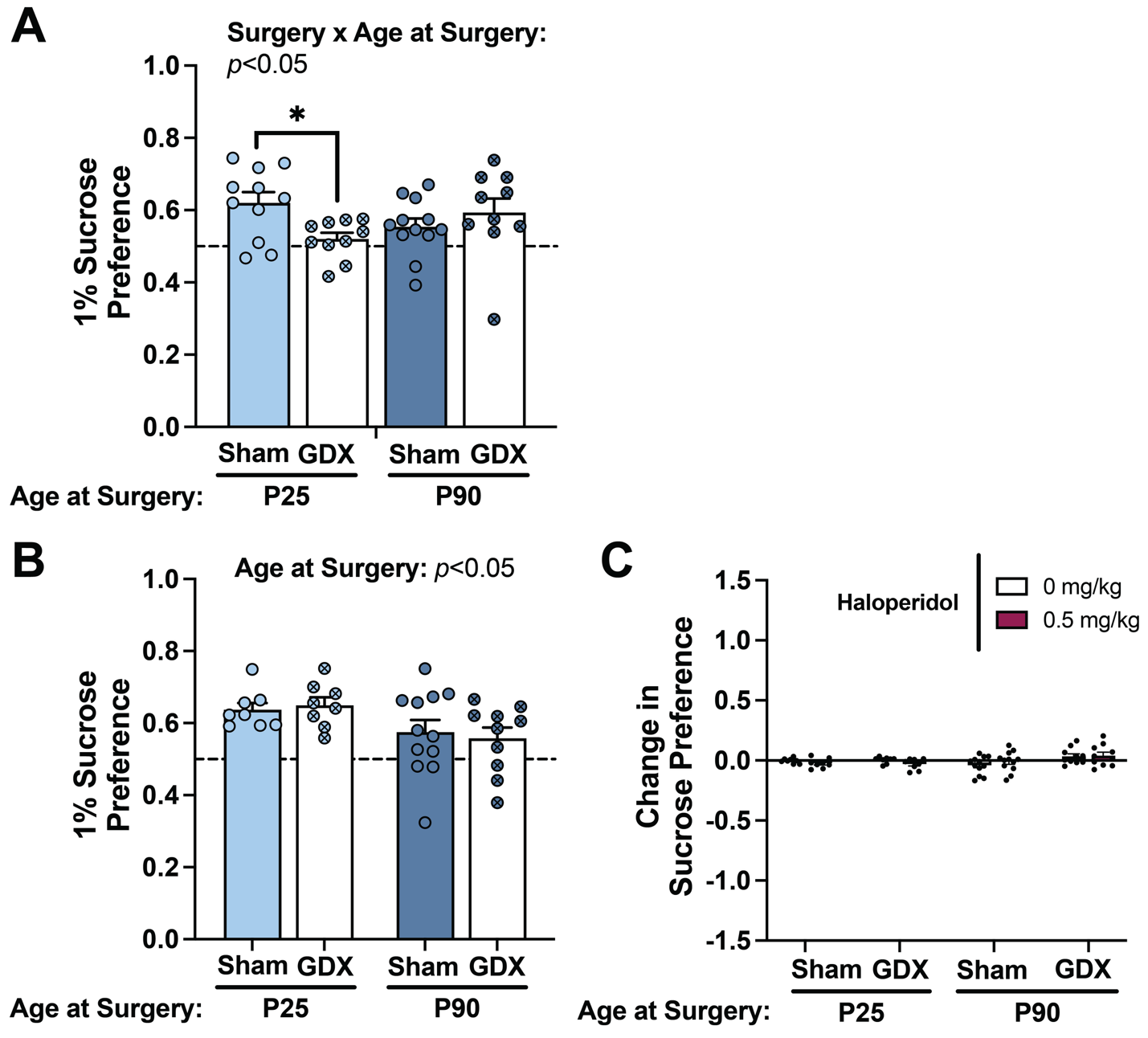

**Supplementary Figure S3.** Baseline sucrose preference testing during PR1-CE (SPT1) and testing during free feeding (SPT3). (**A**) During SPT1, prepubertal, but not postpubertal, ORX resulted in a reduction in sucrose preference relative to prepubertal sham counterparts (simple main effect of surgery with P25 group: **p*<0.05). (**B**) During the baseline of SPT3, younger mice had higher sucrose preference compared to older mice, regardless of surgery (main effect of age: *p*<0.05). (**C**) Haloperidol challenge failed to alter sucrose preference in any groups.
